# Towards the identification and optimization of the “dose-response” relationship of transcranial direct current stimulation

**DOI:** 10.1101/2022.01.22.477353

**Authors:** R. Salvador, M. C. Biagi, M. Pérez Pelegrí, J. Zhou, T. Travison, A. Pascual-Leone, B. Manor, G. Ruffini

## Abstract

Transcranial direct current stimulation (tDCS) is being investigated as a therapeutic tool in several neuropsychiatric disorders, but its mechanisms of action remain incompletely understood and its effectiveness is highly variable from subject to subject. Some of this inter-subject variance is thought to stem from individual differences in head anatomy, which result in different electric field distributions across individuals for the same stimulation settings. In a recent block-randomized, sham-controlled pilot study, 18 older adults with mild-to-moderate motor and cognitive impairments underwent a 2-week, 10-session intervention of 2 mA bipolar tDCS with the anode over the left dorsolateral prefrontal cortex (DLPFC) and cathode over the right supraorbital region (anodal F3 vs. FP2 using sponge electrodes in the 10-20 EEG system). tDCS, compared to sham stimulation, improved average cognitive-motor performance over a two-week follow-up. For a subset of 12 participants from this cohort who underwent structural brain MRIs at baseline, we derived tDCS-induced electric field properties using MRI-derived realistic, individualized finite element modeling. We hypothesized that the electric field component orthogonal (*E_n_*) to the cortex over the left DLPFC would correlate with observed cognitive-motor improvements. Across the 12 subjects the average tDCS-induced electric field over the target area was highly variable with a standard deviation of the surface average *E_n_* over the left DLPFC of 21% of the mean. For those subjects who were randomized to tDCS intervention (n=6), those who exhibited greater improvement in cognitive-motor performance were exposed to greater normal component of the tDCS electric field averaged over the left DLPFC (*r*_95 %_ ∈ [−0.99,−0.59]). In contrast, averaged over the right DLPFC, *E_n_* was a poor predictor of outcome (*r*_95_ % ∈ [−0.08,0.97]), as was the amplitude of the current injected (a widely adopted dosing parameter). In light of these results, we conducted a retrospective modeling experiment using personalized montage optimization algorithms to achieve a desired *E_n_* amplitude over the left DLPFC. The optimized montages induced an average *E_n_* on the left DLPFC comparable to the bipolar montages, but with half the variability (standard deviation was 11% compared to 21% with bipolar montages). In addition, these results suggest that optimized multichannel montages afford more focal targeting, with lower “spill-over” of electric field onto other cortical areas. These proof-of-concept results, though based upon a small sample size, indicate that assessment of the E-field distribution is critical when evaluating the effects of tDCS intervention. They also reinforce the importance of controlled cortical electric field dosing in tDCS and the potential advantages of optimized multichannel stimulation.

Main findings:

- The lDPFC averaged normal component of the electric field correlated with cognitive-motor improvements following tDCS intervention.
- In older adults, head and brain anatomical differences resulted in high inter-subject variability of tDCS-induced electric fields.
- tDCS montages optimized from modeling and delivered via multichannel electrode arrays can control variance and maximize electric field on target, while minimizing effects elsewhere.

## 1. Introduction

Determining the ideal dose parameters – electrode positions and currents, duration and number of sessions, etc. – of transcranial direct current stimulation (tDCS) is fundamental for optimal and reliable therapeutic effectiveness. Currently, however, there are no gold standards for defining these parameters. This limitation stems from 1) insufficient knowledge of the effects of tDCS, including the mechanisms of interaction between the electric field (E-field) induced in the brain by tDCS during stimulation and cortical neurons and their plastic effects, and 2) the lack of an analytic and technical pipeline enabling optimization of dose parameters across individuals. For these reasons, many attempts to study tDCS still rely on bipolar montages that employ large electrodes and affect large portions of the brain (Miranda et al., 2013). In such montages, the anode/cathode is typically placed over the target area (e.g., the primary motor cortex [M1] or the left dorsolateral prefrontal cortex [DLPFC] (Lefaucheur et al., 2017)) in an attempt to increase/decrease its cortical excitability, respectively. The so-called “return electrode”, of opposite polarity to the “active” one, is placed in a region far away located in the opposite hemisphere or even extra-cephalically. Even studies that rely on multichannel stimulation (i.e., more than 2 channels) using smaller electrodes often resort to relatively simple pre-defined montages, such as the 4×1 montage (one central electrode surrounded by 4 electrodes of opposite polarity, (Datta et al., 2009)). Furthermore, the currents employed by these studies are usually kept constant across all subjects (1-2 mA, (Antal et al., 2017). Such “one-model fits all” approaches to montage definition contribute to high inter-subject variability in E-field distribution characteristics, as evaluated in computational modeling studies that employ realistic head models built from individual MRIs (Datta et al., 2012; Laakso et al., 2015). This variability has been linked to underlying anatomical differences between subjects – in particular, the thickness of tissues such as the skull and the cerebrospinal fluid (CSF) (Opitz et al., 2015). These effects are likely increased in older populations, due to varying degrees of brain atrophy and other structural alterations associated with aging.

A complete understanding of how E-field differences relate to the physiologic or functional impact of stimulation outcome must stem from knowledge of the mechanisms behind its interactions with neurons. The effects of tDCS are thought to result in part from the coupling of the electric field to populations of pyramidal cells, leading to coherent changes in membrane potentials (Ruffini et al., 2013; Lefaucheur and Wendling, 2019). Glial cells are also exposed to such effects (Ruohonen and Karhu, 2012). Elongated cells are influenced mostly by the component of the electric field parallel to their trajectory (Bikson et al., 2004; Datta et al., 2008a; Fröhlich and McCormick, 2010; Ranck Jr., 1975; Rattay, 1986; Rushton, 1927), and knowledge about the orientation of the electric field is crucial in predicting the effects of stimulation (Kebakov et al, 2012, Sun et al, 2020). The components of the field perpendicular and parallel to the cortical surface are of special importance since pyramidal cells near the cortical surface are mostly aligned perpendicular to the cortex, while many cortical interneurons and axonal projections of pyramidal cells tend to align parallel to this surface (Day et al., 1989; Fox et al., 2004; Kammer et al., 2007). Glial cells are arranged in similar patterns as neurons. Thus, an important element for modeling tDCS effects is to account for the field distribution and orientation relative to the grey matter (GM) and white matter (WM) surfaces. Since the spatial scale of variation of tDCS electric fields is relatively large (of the order of a cm) compared to the length of neuronal processes in the GM, the above considerations imply that the effect of stimulation should correlate with spatial averages of the component of the electric field normal (*E*_n_) to the cortex over cm scales.

In this study, we tested the hypothesis that the E-field normal to the cortical surface averaged over the target area correlates with the functional changes observed following intervention and explains a large portion of the inter-individual variability. Based on this, we sought to establish the foundations for a targeting approach taking into account the direction of the E-field in the cortex that i) accounts for individual head anatomy and ii) takes advantage of multichannel montages to optimize the match of E-field distribution with the target and minimizes inter-subject variability.

## 2. Materials and methods

### 2.1. Study design – stimulation protocol and performance recording

The data used in this work were collected in a double-blinded, block-randomized, sham-controlled pilot trial of a two-week tDCS intervention (NCT02436915) (Manor et al., 2018). Written informed consent was obtained and all study procedures were approved by the Hebrew Senior Life Institutional Review Board. Eighteen older adults aged 65 years and older who presented with mild-to-moderate motor and cognitive impairments, yet without overt illness or disease, completed baseline assessments of cognitive-motor function. The primary behavioral outcome for that study was the dual task cost (i.e., percent decrement) to gait speed, derived from a dual task paradigm in which participants completed trials of walking both with and without simultaneous performance of a serial subtraction cognitive task.

Participants were randomized to receive a two-week, ten-session tDCS intervention targeting the left DLPFC. Each tDCS session consisted of 20 minutes of continuous stimulation with a maximum 2.0 mA current. Stimulation was delivered with the Starstim system (Neuroelectrics, Barcelona, Spain) connected to saline-soaked synthetic sponge electrodes (SPONSTIM by Neuroelectrics, with 25 cm^2^, circular electrodes) placed over F3 (anode) and FP2 (cathode) of the 10/10 EEG electrode placement system. Follow-up assessments were completed within three days and two weeks after the intervention.

Of the total cohort of 18 study participants, a subset of the 12 (8 women; age range= 66-93 years; average age ± standard deviation = 76±9 years; tDCS/sham arm allocation = 6/6) who were interested and eligible for MRIs also completed a baseline structural brain MRI using a GE Signa HDxt 3 Tesla system with an 8-channel head coil within the Center for Advanced MR Imaging at the Beth Israel Deaconess Medical Center. A T1-weighted MDEFT (Modified Driven Equilibrium Fourier Transform) scan (inversion time=1100ms, TR=6.616ms, TE=2.84ms, flip angle=15°, resolution= 1.000mm × 0.9375 mm × 0.9375 mm) was acquired for whole-brain high-resolution anatomy.

### 2.2. Numerical calculation of E-field in bipolar montages with sponge electrodes

Realistic head models of the twelve participants were created from their respective T1 weighted brain MRIs. The images were segmented into white-matter (WM), grey-matter (GM), cerebrospinal fluid (CSF), air filled sinuses, skull and scalp using FreeSurfer (v6.0, https://surfer.nmr.mgh.harvard.edu/) and SPM 8 (https://www.fil.ion.ucl.ac.uk/spm/software/spm8/) with the MARS package (Huang et al., 2013). FreeSurfer was used to segment the GM and WM (including cerebellar WM and GM) and MARS was used to segment the remaining tissues. The segmentation results were combined using custom Matlab scripts (r2018a, www.mathworks.com) which a) ensured all tissues were surrounded by at least a 1 mm layer of another tissue, b) generated surfaces of all the tissues, and c) smoothed all the tissues and resolved any intersections between surfaces. These procedures were performed using Matlab’s Image Processing Toolbox and Iso2Mesh (Fang and Boas, 2009). Electrode positions on the scalp were defined based upon the 10/10 EEG system (Jurcak et al., 2007) and determined via manual identification of the *nasium*, *inium* and the pre-auricular points. The electrodes used in the bipolar montage stimulation (cylindrical, 2.8 cm radius, 25 cm^2^ area) were then placed in positions F3 and FP2. After the electrodes were placed, finite element volume meshes of the full head (with electrodes) were created using Iso2Mesh. The mesh comprised about 6 million tetrahedra. The geometry of the head, the electrodes and the some of the tissues are shown in **Figure 1** for one of the subjects.

**Figure 1:**
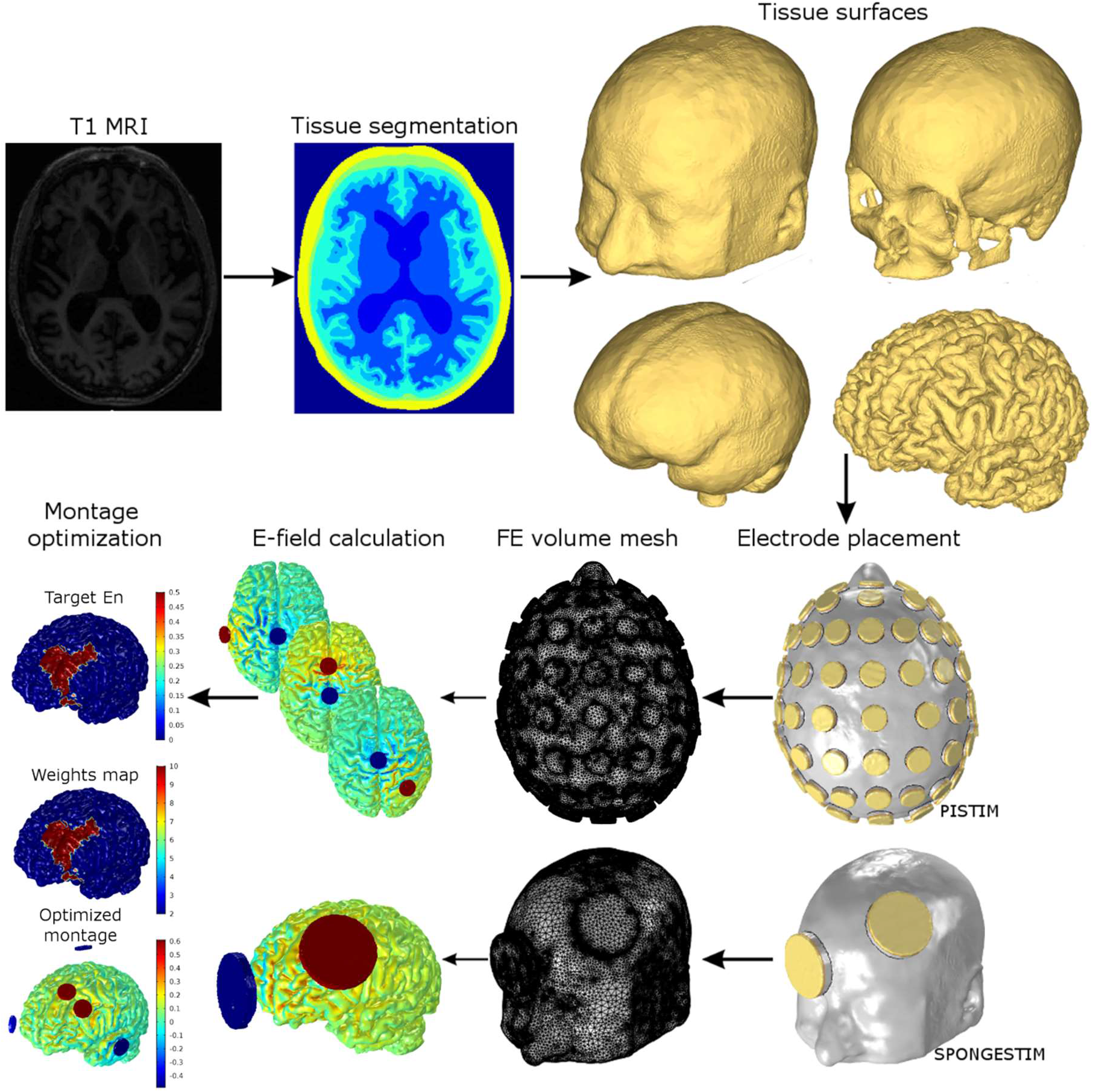
Processing pipeline for every subject in this study. The T1-weighted MRIs were segmented into six tissues using *MARS* and *Freesurfer*. Triangulated surface meshes for each tissue are then created and two types of electrodes are represented in the head model: SPONTIM electrodes (cylindrical, 25 cm^2^, anode over F3 and cathode over FP2) and PISTIM electrodes (cylindrical, 1 cm radius, subset of 61 positions of the 10-10 EEG system). Volume meshes are then created and imported into *Comsol*, where the E-field calculation is performed. For the model with the PISTIM electrodes, 60 E-field calculations are performed (all possible bipolar montages with Cz as a common cathode, −1.0 mA) which allow to calculate the E-field using the principle of linear superposition (see (Miranda et al., 2013, Ruffini et al., 2014)). A map of the left DLPFC region is then created and the montage optimization run with the following parameters: target E_n_ in the left DLPFC: +0.50 V/m (pointing into cortex); weights in target region: 10; target E_n_ in rest of the GM surface: 0.0 V/m; weight in rest of the GM surface: 2; maximum current per electrode: 2.0 mA; maximum total injected current: 2.0 mA; maximum number of electrodes: 6.

Each mesh was imported into *Comsol* (v5.3a, www.comsol.com) in which appropriate tissue conductivities for DC-low frequency range were assigned: 0.33 S/m, 0.008 S/m, 1.79 S/m, 0.40 S/m, 0.15 S/m and 10^-5^ S/m respectively for the scalp, skull, CSF, GM, WM and air (Miranda et al., 2013). All tissues were assigned isotropic conductivity values. The electrodes were represented as conductive media with an isotropic and homogenous conductivity of 4.0 S/m. The voltage difference between the outer surfaces of the electrodes was set so that the injected current was of 2.0 mA (anode over F3, cathode over FP2). Other boundary conditions include continuity of the normal component of the E-field in the inner surfaces of the model and electrical insulation in the outer surface of the model. Laplace’s equation was solved for the electrostatic potential (*V*) using Lagrange second order finite elements. We then calculated the electric field (E-field) by taking the negative spatial gradient of *V*.

The electric field components (normal or orthogonal component of the E-field, *E_n_*, magnitude of the tangential component of the E-field, *E_t_*, and the E-field’s magnitude, *E*) on the GM surface (GM-CSF interface) were calculated for each subject. Furthermore, cortical surface average values of the components of the E-field and norm in the left and right DLPFC were also calculated (hereby represented as 〈*E_n_*〉, 〈*E_t_*〉 and 〈*E*〉). To identify these regions in each subject, we used the BA (Brodmann Area) parcellation of the Colin27 brain template (http://www.bic.mni.mcgill.ca/ServicesAtlases/Colin27) provided by MRICro (http://people.cas.sc.edu/rorden/mricro/). Both the Colin27 template and each of the subjects head models were then put in a common space using *FreeSurfer*’s transformation from RAS space to Tailarach space (https://surfer.nmr.mgh.harvard.edu/fswiki/CoordinateSystems), and the parcellation on a subject’s head was matched to that on the template. The DLPFC regions were then defined as BA 9 and 46 (see **Figure 1** for this mapping on one of the subjects).

### 2.3. Montage optimization

For the optimizations, a version of the head model was created with 64 electrodes (subset of the 10/10 EEG system) representing gelled PISTIM *Ag/AgCl* electrodes (https://www.neuroelectrics.com/products/electrodes/pistim/). The latter were modeled as cylinders (1 cm radius, *π* cm^2^ area, 3 mm thickness) representing the conductive gel under the electrodes, as shown in **Figure 1**. This model was then imported into *Comsol* and the same conductivity values as the ones used in the bipolar montage models were assigned to the tissues/materials. In *Comsol*, the E-field normal to the GM surface (*E_n_*) was calculated for all the bipolar montages with Cz as a cathode (−1 mA) and each of the other electrodes as the anode (+1 mA). Linear combinations of these distributions allow the calculations of the *E_n_* field distribution with any montage employing these electrodes (Ruffini et al., 2014). The distributions were used as the input to the optimization algorithm.

Montage optimizations were performed using the *Stimweaver* algorithm, described in (Ruffini et al., 2014). This algorithm optimizes for the *E_n_* component of the E-field, considering that an E-field directed towards the cortical surface and normal to the interface (positive *E_n_* in the convention followed in this study) will depolarize the soma or axon terminals of pyramidal cells, with the opposite happening for an *E_n_* field directed out of the cortical surface (negative *E_n_*). In this optimization algorithm, the optimal montage (electrode positions and currents) is considered the one that minimizes the weighted least squares difference between the target *E_n_* and the *E_n_* induced by the montage. The weights (integers between 2 and 10) and target *E_n_* (+0.25 V/m for excitatory effects, −0.25 V/m for inhibitory effects and 0 V/m for no effects) are assigned to all nodes of the cortical surface mesh. For each optimization, a target *E_n_* of +0.25 /m was assigned to the left DLPFC, with maximum weight, as shown in **Figure 1**. The rest of the cortical areas were assigned to a target *E_n_* of 0 V/m with a low weight (w=2). The optimization was constrained for the maximum current per electrode 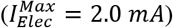 and total injected current (the sum of all the positive currents, 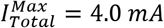). A genetic algorithm was employed to find the solution with a maximum number of 6 electrodes. The surface averages of the components of the Efield in the left and right DLPFC were also calculated for each optimized montage.

### 2.4. Statistical analysis

This was an exploratory study of the statistical relationship between electric field and response (see section 3.2). Reasonableness of assumption of linear association between variables was ascertained using scatterplot smoothing. Following this, linear regressions were calculated using the *statsmodel* (Seabold and Perktold, 2010) library in Python. Specifically, we used the *ols* (ordinary least squares) function to generate *r* values and associated 95% confidence intervals. This methodology was also applied to calculate the statistical significance of correlations between anatomical properties and average E-field values (see section 3.4). Comparisons between surface average values of the different E-field components between the bipolar and multichannel montages (see section 3.3) were calculated with Welch’s t-test (two-tailed), as implemented by the *statnnot* library in Python (https://github.com/webermarcolivier/statannot).

## 3. Results

### 3.1. E-field distribution analysis: bipolar montage with sponge electrodes

The distribution of the different components of the E-field induced during the bipolar montage is shown in ***Figure 2*** for the subjects with highest (subject #4) and lowest (subject #7) average normal field 〈*E_n_*〉, respectively in the top and bottom rows. The distribution of the normal component of the E-field (***Figure 2a/d***) displays the expected pattern, with high E-field values below the electrodes and in areas in between the electrodes. Both positive and negative *E_n_* values are found below both electrodes, depending on the direction of the field with respect to the cortical folds. The mean < *E_n_* > across all subjects is positive for the left DLPFC (average and standard deviation of 0.048±0.010 V/m) and negative for the right DLPFC (−0.019±0.007 V/m). The other components of the E-field (tangential and magnitude, as shown in ***Figures 2b/e*** and ***c/f***) yield higher surface average values and are more symmetric across hemispheres: < *E_t_* >_*left DLPFC*_ = 0.14 ± 0.04 *V*/*m*, < *E_t_* >_*rght DLPFC*_ = 0.13 ± 0.02 *V*/*m*. < *E* >;_*left DLPFC*_ = 0.21 ± 0.05 *V*/*m*. < *E* >_*right DLPFC*_ = 0.19 ± 0.03 *V*/*m*. The values of 〈*E_n_*〉. 〈*E_t_*〉 and 〈*E*〉 for both areas and for all the subjects are presented as Supplementary material (***Table s1***).

**Figure 2:**
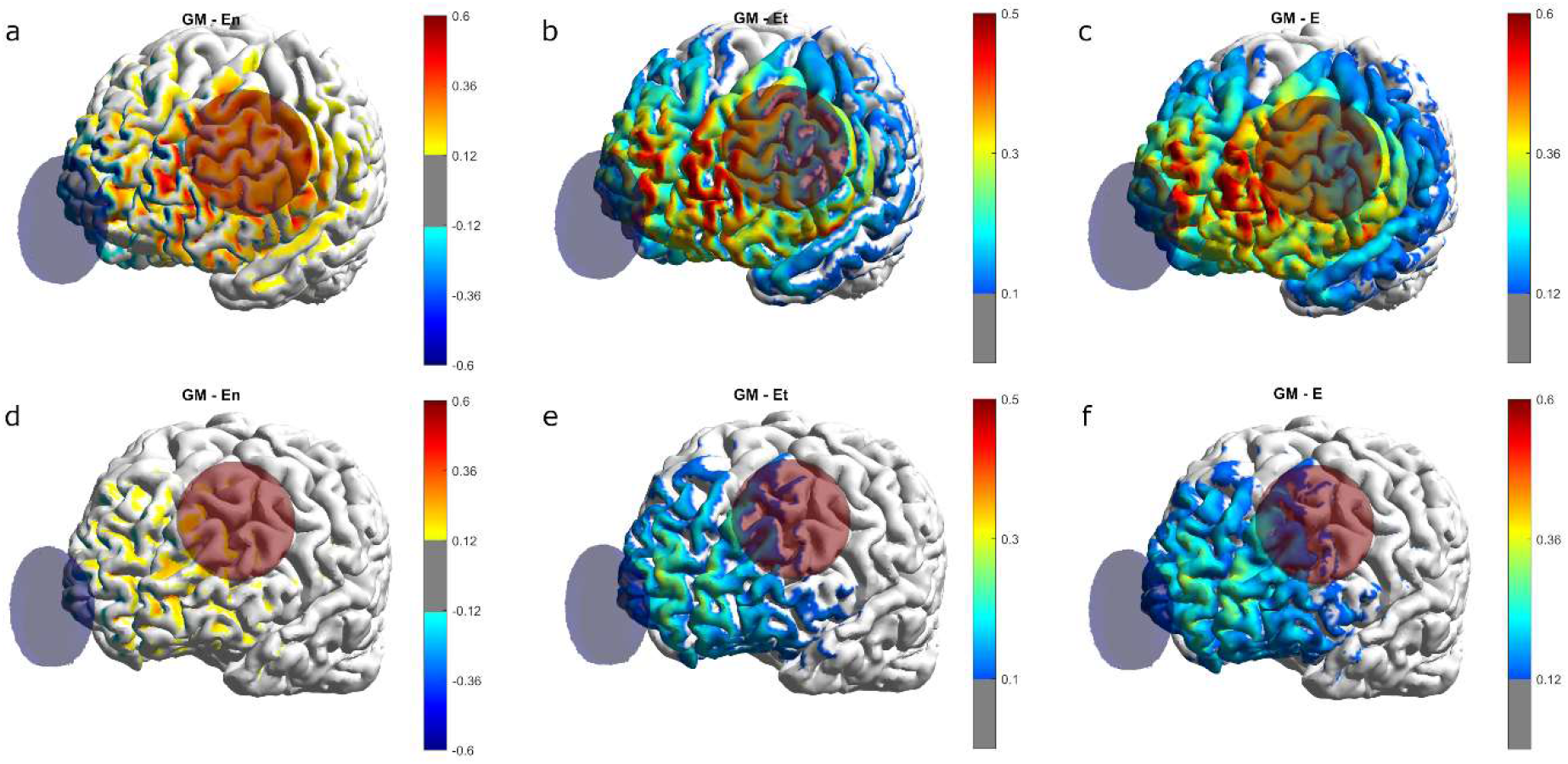
E-field distribution in the bipolar montage for the subject with highest (top row, a-c) and lowest (bottom row, d-f) 〈*E_n_*〉_*left DLPFC*_. The top 1^st^ column (a) displays the distribution of *E_n_*; the 2^nd^ column (b) the distribution of *E_t_* and the 3^rd^ (c) the E-field’s magnitude. All values (in V/m) are plotted using a common scale across montages (but different for each E-field component) which with a threshold of 20% of the maximum value. Anodes are shown in red and cathodes in blue.

During the stimulation protocol of the initial study, the injected current was set to a lower value for some of the subjects due to complaints of burning/itching sensations under the electrode. This has the effect of scaling down the E-field values (and averages values) proportionally to the reduced amount, as there is a linear relation between the current and the induced E-field. ***Table s1*** also presents the scaled down average values for all the subjects to which this applies.

### 3.2. Correlation of electric field with impact on cognitive-motor performance

For the six subjects who were randomized to tDCS intervention, we calculated a linear fit between the subject-level average E-field components over both left and right DLPFC and the pre-to-post intervention change in cognitive-motor performance. Cognitive-motor performance was defined as the dual task cost to gait speed; that is, the percent reduction in gait speed when walking and performing verbalized serial subtractions, as compared to walking alone (see ***Figure 3***). Linear regression showed that those subjects who received greater < *E_n_* > over the putative target area, < *E_n_* >_*left DLPFC*_, exhibited greater pre-to-post reduction (i.e., improvement) in dual task cost (Pearson’s *r* 95% confidence interval: [−0.99, −0.59]). Similar levels of correlation were observed with < *E_t_* >_*left DLPFC*_ (*r* ∈ [−0.99,−0.59]) and < *E_t_* >_*right DLPFC*_ (*r* ∈ [−0.99,−0.56]). The only non-significant fit was that of the normal component of the E-field in the right DLPFC: < *E_n_* >_*rlght DLPFC*_ (*r* ∈ [-0.08,0.97]). Since the actual currents employed in the bipolar stimulation were reduced for some of the subjects from the initially planned 2.0 mA, we also calculated a linear regression between the total injected current (here defined as the sum of all the positive currents) and performance change (not shown in the figure). The resultant fit was found to be non-significant (*r* ∈ [-0.96,0.30]).

**Figure 3:**
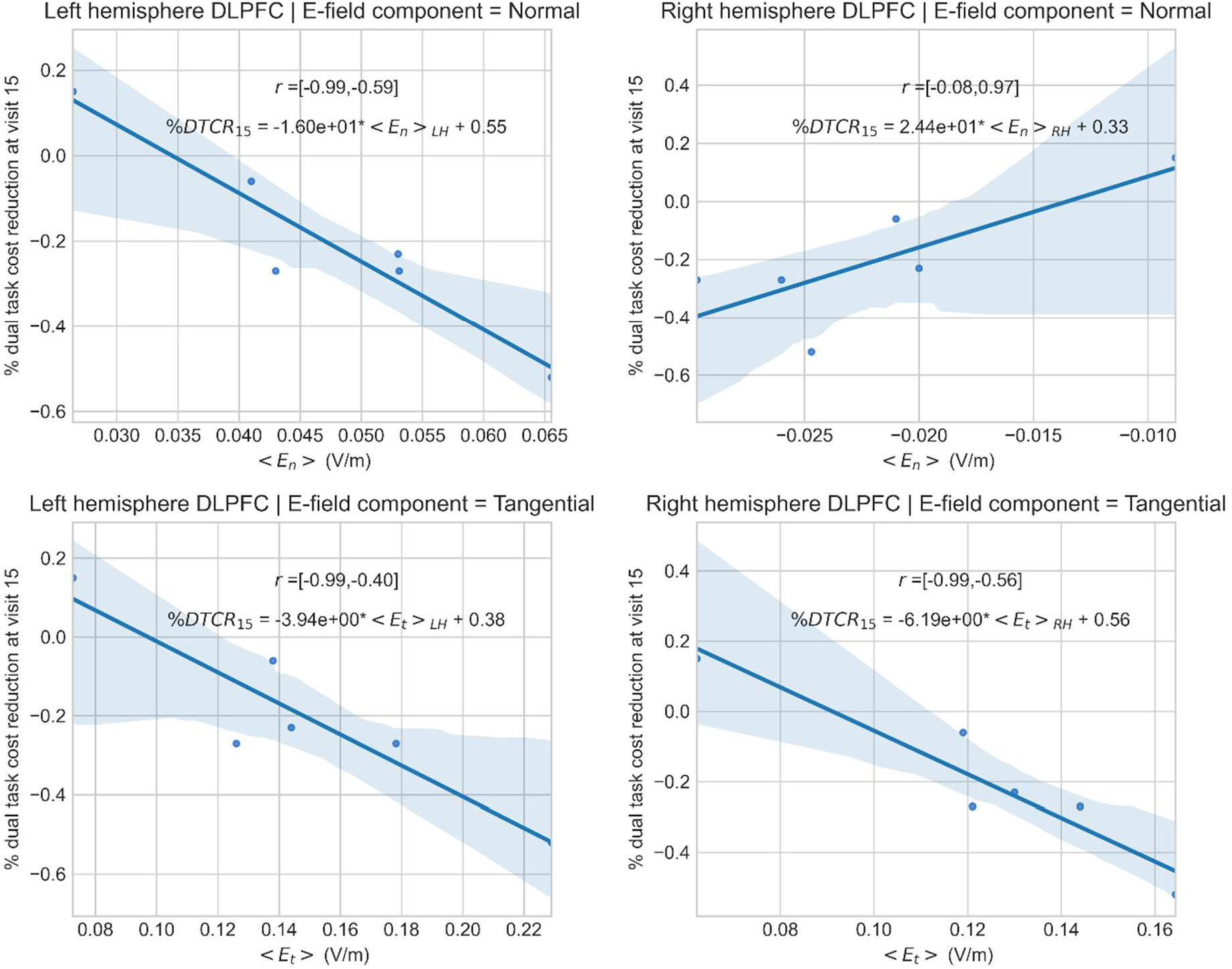
Dual-task cost reduction (at visit 15) as a function of <*E_n_*> and <*E_t_*> for both areas of interest (left DLPFC and right DLPFC). The linear fits are presented for each. Data were collected for 6 of the subjects in this study. The shaded areas represent 95% confidence intervals to the fit, generated by bootstrapping. 95% confidence intervals for Pearson’s correlation coefficient are also shown, as well as the expression for the linear fit.

### 3.3. E-field distribution analysis: optimized montage with multichannel high resolution tDCS

Motivated by the above results, we sought to investigate the possible advantages of using a montage optimization approach targeting one specific region (left DLPFC) and optimizing for one component of the Efield (*E_n_*). The optimized montage for most subjects placed three anodes (out of a maximum of six) over the left DLPFC (AF7, FC3, F1, F3 and/or AFz) summing a total current that varied between 2.7 and 4.0 mA, surrounded by cathodes (usually located over Fp1, FT7, Fpz, FC5 and/or Cz) (see ***Figure 4a/d***). ***Table s2*** in supplementary material provides the electrode positions and currents in each optimized montage. This montage results in a focal *E_n_*-field, with much higher surface averages over the left DLPFC than the right DLPFC: < *E_n_* >_*left DLPFC*_ 0.047 ± 0.005 *V*/*m* vs. < *E_n_* >_*rightDLPFC*_= 0.009 ± 0.005 *V*/*m*. Surface averages for the other E-field components are also higher in the left DLPFC than in the right area, but the relative difference is smaller: < *E_t_* >_*left DLPFC*_= 0.088 ± 0.004 *V*/*m*, < *E_t_* >_*right DLPFC*_= 0.052 ± 0.010 *V*/*m*, < *E* >_*left DLPFC*_= 0.128 ± 0.006 *V*/*m*, < *E* >_*right DLPFC*_= 0.074 ± 0.013 *V*/*m*. ***Figure 5*** provides the comparison of the surface average values for the bipolar and multichannel optimized montages, averaged across all subjects. The values of 〈*E_n_*〉. 〈*E_t_*〉 and 〈*E*〉 for each subject in the optimized montages are presented in ***Table s1***.

**Figure 4:**
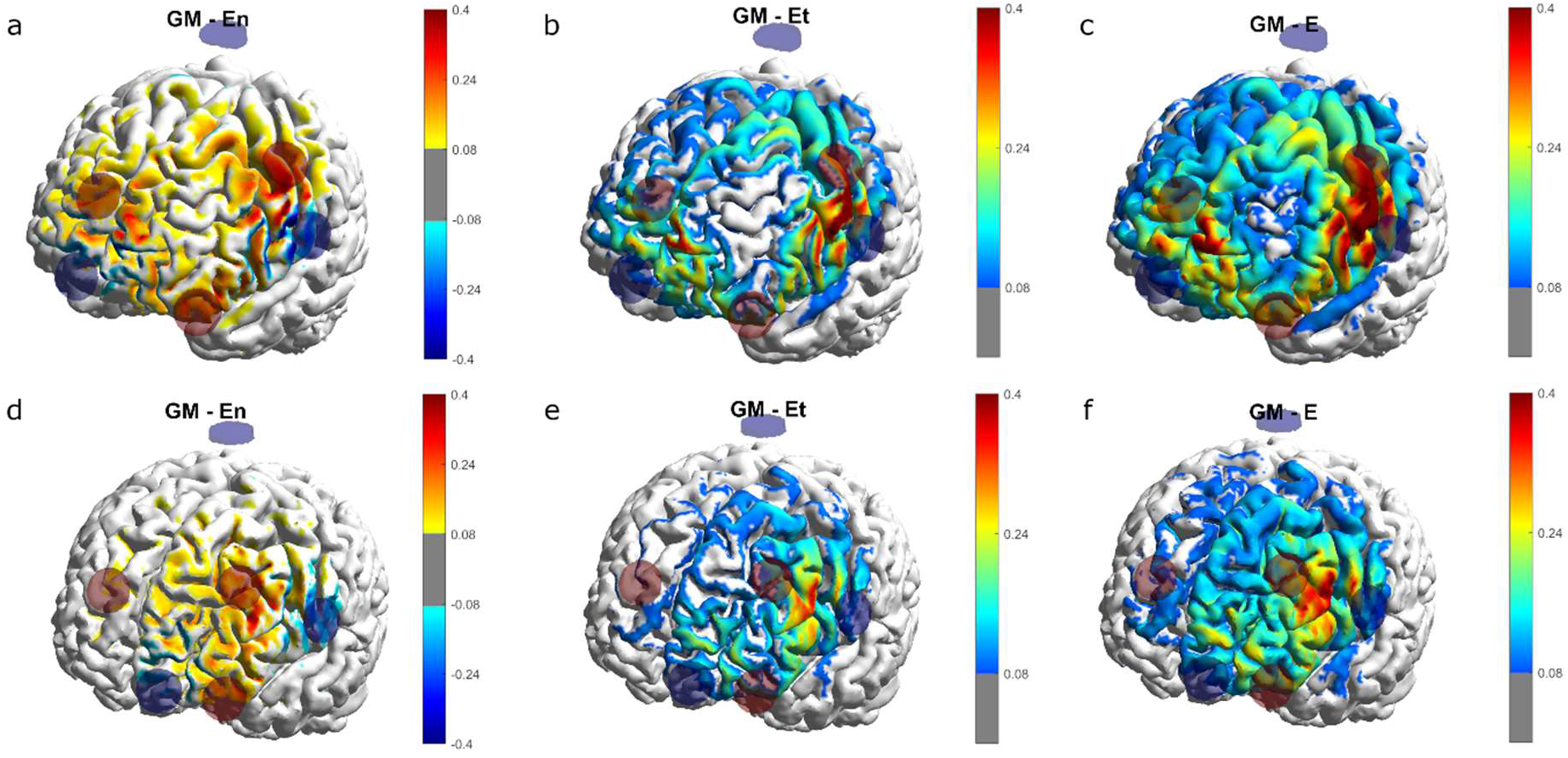
E-field distribution in the optimized montage for the subjects with highest (top row) and lowest (bottom row) 〈*E_n_*〉_*left DLPFC*_. The 1^st^ column shows the distribution of *E_n_*; the 2^nd^ column the distribution of *E_t_* and the 3^rd^ the E-field’s magnitude. All values (in V/m) are plotted using a common scale across montages (but different for each E-field component) with a threshold of 20% of the maximum value. Anodes are shown in red and cathodes in blue.

**Figure 5:**
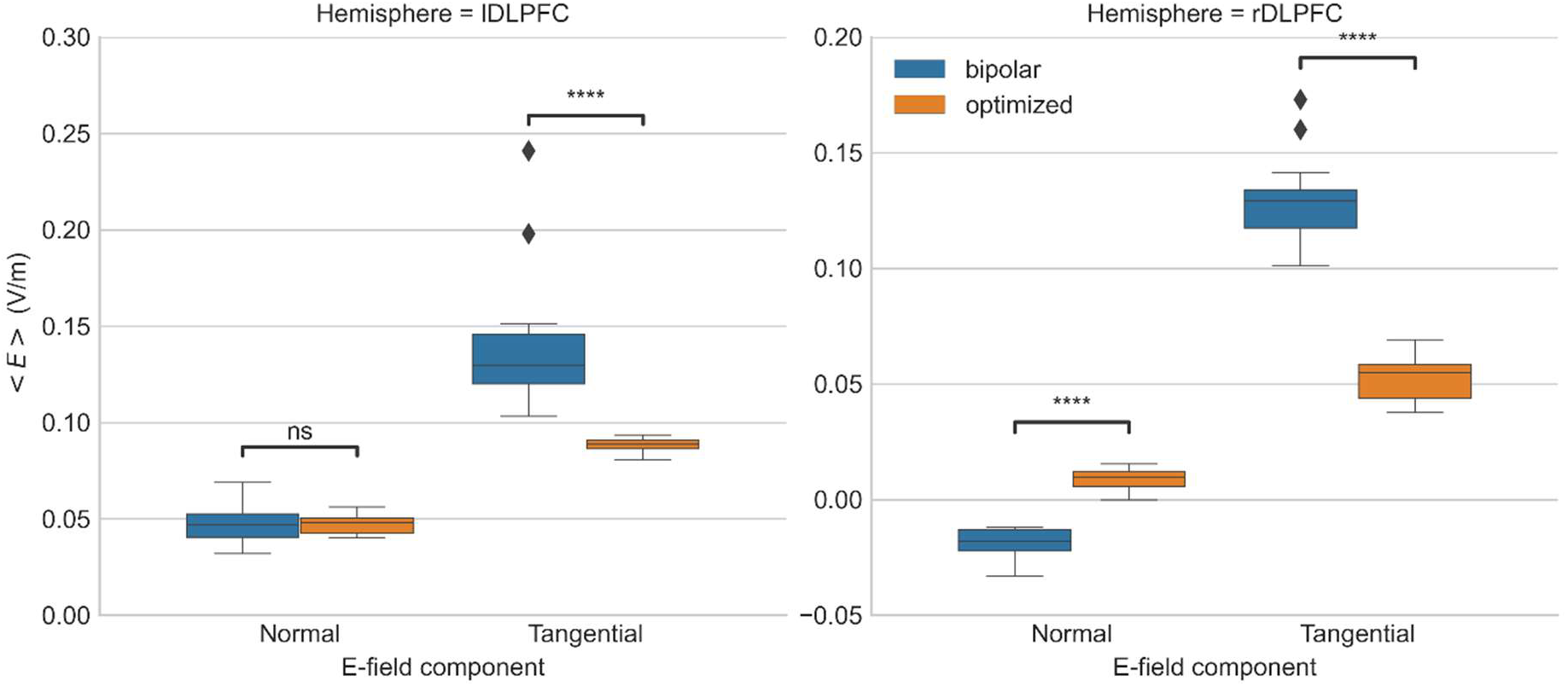
Box and whisker plots of the surface average values of all E-field components for bipolar and optimized multichannel montages across all subjects (N=12). Statistical significance of the difference between the means of the average E-field between montages is also shown in the plots for all E-field components and for both areas of interest (left DLPFC and right DLPFC). Statistical significance was tested using Welch’s t-test. Legend: *ns*: Not significant (*p* > 0.05), ****: *p* < 10^-4^, ♦: outlier.

### 3.4. Drivers of intersubject average electric field variability

To determine the influence of tissue anatomy in the E-field distribution induced with both types of montages studied in this paper, we calculated the correlation between the average E-field components and the inverse of the sum of the volumes of the CSF, skull and scalp tissues. The linear fits to the data for the bipolar/optimized montages are shown in ***figure 6***. Correlations of < *E_n_* > and < *E_t_* > over both the left and right DLPFC were statistically significant for the bipolar montage. For the optimized montages, < *E_n_* > correlated significantly with the inverse of the sum of the volumes, but not < *E_t_* >.

**Figure 6:**
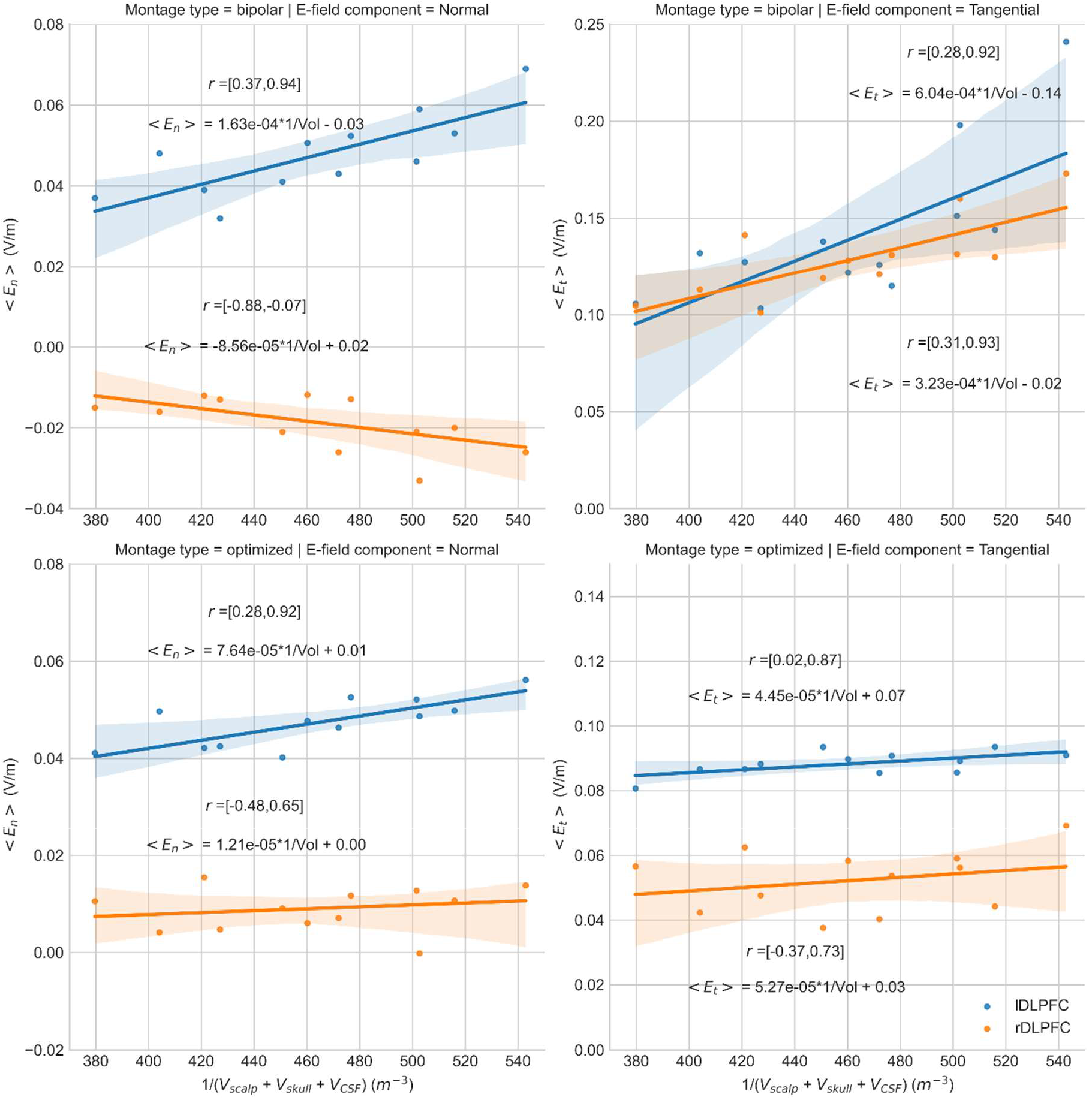
Surface average values of all E-field components (from left to right: <*E_n_*> and <*E_t_*>) for all subjects in both the left DLPFC (blue) and right DLPFC (orange) as a function of the inverse of the sum of the total volumes of the scalp, skull and CSF tissues for the bipolar (top row) and optimized (bottom row) montages. The shaded areas represent 95% confidence intervals to the fit, generated by bootstrapping. 95% confidence intervals for Pearson’s correlation coefficient are also shown, as well as the expression for the linear fit.

## 4. Discussion

In this paper we provide proof of concept that the behavioral and cognitive impact of tDCS intervention depends in part upon the characteristics of the E-field generated in the target region for stimulation. Specifically, analyzing data from a previously published pilot randomized controlled trial (RCT) (Manor et al., 2018), we observed that participants in whom the normal component of the tDCS-induced E-field averaged over the left DLPFC cortical area was greater, exhibited greater functional improvement. We also observed that a “one-model-fits-all” stimulation protocol leads to substantial inter-subject variability in the surface average values of all E-field components, even when considering a similar total current across subjects (2.0 mA). Linear regression models predicted that this variability in E-field explained 61% to 89% of the inter-subject variability in outcome (depending on the E-field component that was considered). The variability in average values of the different E-field components in the DLPFC in both hemispheres could be explained in part with variability of anatomic features, here quantified as the inverse of the sum of the volumes of the skull, scalp, and CSF. These results emphasize that the practice of reporting dose parameters only (currents, electrode type and positions) is insufficient to properly predict the outcome of stimulation. Also, keeping these parameters fixed across all subjects does not guarantee a homogeneous E-field distribution in the brain, which partly explains the intersubject variability in intervention effectiveness.

### 4.1. Relevant components of the E-field

Based on *in-vitro* and animal work (Ruffini et al., 2013), it is usually assumed that the cortical normal component of the E-field to the cortical surface is the most influential to modulate cortical excitability, which in turn leads to changes in functional outcome. Furthermore, many of the results presented focus on calculations of surface average values of the E-field components. Correlation with intervention-induced changes in functional performance for those six subjects who were randomized to active tDCS arm supports our initial hypothesis, as the coefficient of determination for the linear fit between < *E_n_* >_*left DLPFC*_ and the change in cognitive-motor performance was higher (*r* ∈ [−0.99, −0.59]) compared to other test E-Field characteristics. Furthermore, the coefficient of determination for the fit of < *E_n_* >_*right DLPFC*_ was the lowest (*r* ∈ [−0.08,0.97]), indicating that the average value of *E_n_* over the left DLPFC was driving the behavioral effects.

Further inspection, however, make the interpretation less clear as the coefficient of determination of < *E_t_* > in both regions is also quite high and statistically significant. This is, however, not surprising, since the components of the electric field are subject to Poisson’s equation, which establishes correlations among them, the montage was the same across subjects and current polarity was not reversed. However, we can further observe that of the fits with high statistical significance, the one with < *E_n_* >_*left DLPFC*_ presents the greatest negative slope. The trend is the opposite on the other hemisphere, but not statistically significant. Furthermore, the putative role of any undirected component in tDCS (tangential component or norm) is doubtful, as it is well known that tDCS effects typically change direction when current direction is reversed, e.g., from anodal to cathodal stimulation (Nitsche et al, 2000). The normal component of the electric field is signed and follows this direction reversal, but the tangential component or the electric field magnitude do not. Thus, we expect that adding a few study subjects with a cathodal stimulation condition would disentangle the role of normal component, tangential and magnitude of the electric field. Future studies should include both directions of the average normal electric field.

While features of the cortical electric field over the target area were highly correlated with observed benefit of the intervention, the total injected current did not predict outcome well. This expected result suggests that the effects of tDCS are cortical rather than peripheral, since current intensity and associated electric fields on the scalp under the electrodes will affect directly the peripheral nervous system outside the skull with little anatomical variance. While these results at least suggest that < *E_n_* >_*leftt DLPFC*_ correlate highly with performance in this particular application, they do not rule out other components of the field to play a role or that other areas to be more relevant – further studies should clarify this.

### 4.2. Intersubject variability and optimization

As mentioned before, our modeling results of the subjects that participated in the RCT showed a high intersubject variability in the E-field distribution. This result is supported by previous studies examining intersubject variability in the E-field distribution in tDCS (Laakso et al., 2015). Based upon these results, we employed a montage optimization algorithm focused on optimizing the normal component of the E-field, as done in other studies (Dagan et al., 2018; Fischer et al., 2017; Ruffini et al., 2014). The individually-tailored montages obtained with this method would have decreased the variability in < ***E_n_*** >**_*left DLPFCffc*_**: standard deviation was 11% of the mean across subjects, compared to 21% with bipolar montages. Since the variability of < ***E_n_*** >**_*left DLPFC*_** significantly correlated with the variability of the functional outcomes of stimulation, this indicates that optimized approaches to montage design can help maintain a more uniform response to stimulation. However, even in this optimized approach the correlation of < ***E_n_*** >**_*left DLPFC*_** with the inverse of the sum of the volumes of non-brain tissues was still statistically significant. It is likely that increasing the maximum available total injected current and/or decreasing the target-*E_n_* field would result in less variability in the average *E_n_* across subjects and, therefore, reduce the correlation with anatomical features. The same would occur with a smaller target area, for which it would be easier to achieve a stable average *E_n_*-field. The optimization does, however, decouple the average *E_n_*-field values from anatomical parameters in the right hemisphere. The same applies for <*E_t_*> in the right hemisphere. This indicates that the optimized montage achieves a more focal E-field located in the left hemisphere as compared to the bipolar montages and, at the same time, decreases the overall contribution of the tangential component of the E-field to the total E-field.

While the increase in focality observed with multichannel optimized montages is not guaranteed to lead to better functional outcome in stimulation (e.g., if the relevant target was not correctly identified), it is an important proof-of-concept towards the design more controllable experiments to further assess the importance of specific cortical regions/E-field components. It should be noted that the optimization approach in (Ruffini2014) can be used to optimize for targets using the tangential component (or magnitude) of the E-field, in case these components do have a role in determining functional outcome (an hypothesis we could not discard completely in this study). It should also be mentioned that it is possible to tune the optimization parameters towards higher average E-field on target. This can be done by increasing the target *E_n_*-field or decreasing the weights in the off-target areas (Salvador et al, 2021). This would come at a cost of worse focality but may result in stronger neuromodulatory effects.

The issue of the influence of personalization of montages in the correlation of the E-field with anatomical factors is crucial in studies with heterogeneous populations or studies with populations that may have anatomical characteristics that affect the E-field distribution. Brain atrophy in an elderly population, for instance, may lead to lower average E-field values on average, as compared to younger populations (for the same current values). This may be offset by employing montage optimization algorithms.

The impact of intersubject variability on the average values of the E-field components can be theoretically dismissed if modeling of the montages is performed before the experiment is conducted and designed to stabilize the average electric field over the target. This can easily be achieved by taking advantage of the linearity between the E-field and the current, and by scaling the currents in each montage so that the average Efield values are the same across subjects. In this scenario, the optimized multichannel montages offer additional advantages over the bipolar montages by allowing for a more focal *E_n_*-field distribution and reducing the average values of the other components of the E-field. However, such an approach may be impractical in some experimental designs where the anatomical MRIs are collected gradually throughout the study – which would make modeling of all subject beforehand impossible. A more practical approach would be to scale the results to an average value calculated a priori on a population with similar demographics or to conduct a group-level montage optimization on the same population (Salvador et al, 2021b).

Another important advantage of optimized multichannel montages is related with tolerability of stimulation. In the original experiment, subject complaints resulted in a decrease of the injected current for some subjects. This further exacerbated intersubject variability in the results. Optimized montages may overcome this issue by reducing the maximum allowed current per electrode (but keeping the total injected current equal). For instance, reducing the maximum current per electrode to 1.0 mA while leaving the maximum total injected current unchanged only resulted on a 5% penalty on < *E_n_* >_*left DLPFC*_ (data not shown) in the optimizations we performed in this study.

### 4.3. Comparison with previous studies

The results presented here are consistent with recent studies that have demonstrated relationships between Efield dose and effects of tES on stimulation in cortical excitability changes of the M1 area following stimulation (Laakso et al, 2019 and Mosayebi-Samani, 2021), on neurotransmitter concentration changes and sensorimotor network connection strength (Antonenko et al, 2021). Our data further emphasizes the need of numerical modeling in experimental planning and assessment of the results of transcranial brain stimulation.

### 4.4. Limitations

The current results are based upon a relatively small number of subjects using the same montage polarity. Although twelve subjects were available for analysis of electric field variability from MRI, only six were allocated to receive the active tDCS intervention. Moreover, we used here *a-posteriori* analysis of data using fixed bipolar montages, which produce quite unspecific electric fields spatially, and induce strong correlations between the E-field characteristics over different cortical areas.

The definition of the left DLPFC area is another parameter that presumably influenced our results. In the current study, the target area was defined by cytoarchitectural data (Brodmann brain parcellation). The left DLPFC, however, may be better defined functionally – leading to a different target area (Cieslik et al., 2013; Mueller et al., 2013). Mapping the target region individually for each subject also seems to be beneficial in the case of the DLPFC, given the high inter-subject variability in resting state functional connectivity reported for this region (Wang et al., 2015). However, it is expected that the functionally defined left DLPFC region is mostly encompassed by the mask defined for each subject. One alternative to this analysis would be to use statistical methods that do not assume any relevant target area, as done in (Laakso et al., 2019). However, our 6-subject sample lacks enough power to draw any reliable conclusions as to the most significant areas.

From a modeling perspective, we utilized head models that included a single compartment for the skull, as opposed to separating between the spongy and compact bone tissues. This was due to the lack of T2 or CT scans for each of the subjects, which made it impossible to segment between these two tissues. The models also considered the GM and WM to have an isotropic conductivity profile, because no diffusion weighted MRI data was acquired. Previous modeling studies (Opitz et al., 2015) have shown that the presence of a spongy bone compartment can lead to significant changes in E-field distribution, as spongy bone is considerably more conductive than compact bone. The lack of an anisotropic model for the conductivity of the brain tissues is expected not to affect the cortical E-field distribution significantly (Salvador et al., 2015). These simplifications apply to all head models, so while the average E-field values and the optimized montage outputs reported here will be changed in more complex models, it is unlikely that they affect the overall correlations with performance metrics.

The manner in which the electrodes are modelled also influences the E-field distribution considerably, depending on electrode type (Saturnino et al., 2015). In this work, sponge electrodes were modeled as a single homogeneous compartment. The actual electrodes comprise a central cylinder of conductive rubber located on top of a saline soaked sponge electrode. A more accurate model for these electrodes was tested against the simplified models used in this study, and it was found to have a very small effect on the E-field distribution (see supplementary material, section 3). This is likely due to the fact that the rubber cylinder insert is only slightly smaller, in diameter, than the saline soaked sponge. A further limitation is the lack of cross-validation of the electrode positions in the actual experiment and those in the head models. This can lead to important changes in the E-field distribution (Opitz et al ??). This cross-validation can be implemented in some studies using MRI acquisitions with the electrodes, as was done in (Antonenko et al, 2019).

## 5. Conclusion

The effects of tDCS are understood to derive from the interaction of the induced electric field with neuronal populations in the cortex. It is possible today to model with reasonable precision such electric fields at the individual level. Thanks to this, it is possible to plan and analyze tDCS interventions in cortical electric field space rather than “electrode space” and current space, much as it has happened in the field of electroencephalography (EEG), where modern practice has shifted from the analysis of data in voltage and electrode space to cortical dipole source space. The present study addresses unknowns about the parameters of the E-field that ultimately determine the impact of tDCS intervention. Our results suggest a possible direct link between tDCS effects in electric field space and the functional impact of a multi-session tDCS intervention. They also point the way to further studies to better dissect the causal contributions of different E-field components and regions in the cortex. For this purpose, model-driven multichannel stimulation provides a promising tool, as it allows to manipulate the spatial distribution of the electric field in the cortex. In addition, we note that studies including both cathodal and anodal conditions can help disentangle the role of signed and unsigned functions of the electric field vector such as the normal component (signed) vs. the tangential component or magnitude (unsigned). Finally, in this work we also provide evidence for how montage optimization may be used to determine dose parameters that minimize the variability in the E-field distribution resulting from anatomical differences across subjects. Future work should extend this analysis by replication in a larger cohort, as well as using optimized montages from the beginning to plan stimulation on an individual level.

## Supplementary material

### 1. Values of average components of the E-field in the two target regions

Table s1 provides the values of the average components of the E-field in both the left DLPFC and the right DLPFC for all the subjects in this study. The results are presented for both the bipolar and optimized montages. For the subjects that were stimulated with lower current, we also present the scaled values of the E-field averages (bipolar montage only).

### 2. Optimized montages

The positions and currents of all the electrodes in the optimized montages are presented in table s2. The table also shows the maximum current per electrode and the total injected current for all montages.

**Table s1:** Surface average values of each component of the E-field (V/m) in the left and right DLPFC for all the subjects in the study. The values were calculated for both the bipolar (BM) and optimized montages (OM). Subjects with an asterisk (*) were in the active stimulation branch of the study. Subjects 4, 5 and 6 were stimulated with a current lower than 2.0 mA. For these subjects, the scaled values of the surface average components of the E-field, for a current of 2.0 mA, are presented as well (top row).

| Subj | $\langle E_n \rangle$ | | | | $\langle E_t \rangle$ | | | | $\langle E \rangle$ | | | |
| --- | --- | --- | --- | --- | --- | --- | --- | --- | --- | --- | --- | --- |
|  | Left DLPFC |  | Right DLPFC |  | Left DLPFC |  | Right DLPFC |  | Left DLPFC |  | Right DLPFC |  |
|  | BM | OM | BM | OM | BM | OM | BM | OM | BM | OM | BM | OM |
| 1* | 0.053 | 0.050 | -0.020 | 0.011 | 0.144 | 0.094 | 0.130 | 0.044 | 0.224 | 0.138 | 0.200 | 0.064 |
| 2* | 0.041 | 0.040 | -0.021 | 0.009 | 0.138 | 0.094 | 0.119 | 0.038 | 0.203 | 0.132 | 0.179 | 0.057 |
| 3* | 0.043 | 0.046 | -0.026 | 0.007 | 0.126 | 0.085 | 0.121 | 0.040 | 0.181 | 0.123 | 0.180 | 0.057 |
| 4* | 0.069 | 0.056 | -0.026 | 0.014 | 0.241 | 0.091 | 0.173 | 0.069 | 0.326 | 0.133 | 0.252 | 0.097 |
| (1.9 mA) | 0.066 |  | -0.025 |  | 0.229 |  | 0.164 |  | 0.310 |  | 0.239 |  |
| 5* | 0.048 | 0.050 | -0.016 | 0.004 | 0.132 | 0.087 | 0.113 | 0.042 | 0.213 | 0.132 | 0.168 | 0.063 |
| (1.1 mA) | 0.026 |  | -0.009 |  | 0.072 |  | 0.062 |  | 0.117 |  | 0.093 |  |
| 6* | 0.059 | 0.049 | -0.033 | -0.000 | 0.198 | 0.089 | 0.160 | 0.056 | 0.285 | 0.131 | 0.238 | 0.079 |
| (1.8 mA) | 0.053 |  | -0.030 |  | 0.178 |  | 0.144 |  | 0.257 |  | 0.215 |  |
| 7 | 0.032 | 0.043 | -0.013 | 0.005 | 0.103 | 0.088 | 0.101 | 0.048 | 0.142 | 0.123 | 0.149 | 0.068 |
| 8 | 0.039 | 0.042 | -0.012 | 0.015 | 0.127 | 0.087 | 0.141 | 0.062 | 0.194 | 0.122 | 0.205 | 0.091 |
| 9 | 0.037 | 0.041 | -0.015 | 0.011 | 0.106 | 0.081 | 0.105 | 0.057 | 0.162 | 0.120 | 0.162 | 0.079 |
| 10 | 0.046 | 0.052 | -0.021 | 0.013 | 0.151 | 0.086 | 0.132 | 0.059 | 0.215 | 0.126 | 0.194 | 0.080 |
| 11 | 0.051 | 0.048 | -0.012 | 0.006 | 0.122 | 0.090 | 0.128 | 0.058 | 0.170 | 0.126 | 0.191 | 0.081 |
| 12 | 0.052 | 0.053 | -0.013 | 0.012 | 0.115 | 0.091 | 0.131 | 0.054 | 0.175 | 0.136 | 0.184 | 0.078 |
| $\mu$ | 0.048 | 0.047 | -0.019 | 0.009 | 0.14 | 0.088 | 0.13 | 0.052 | 0.21 | 0.128 | 0.19 | 0.074 |
| $\pm \sigma$ | $\pm 0.010$ | $\pm 0.005$ | $\pm 0.007$ | $\pm 0.005$ | $\pm 0.04$ | $\pm 0.004$ | $\pm 0.02$ | $\pm 0.010$ | $\pm 0.05$ | $\pm 0.006$ | $\pm 0.03$ | $\pm 0.013$ |

**Table s2:**
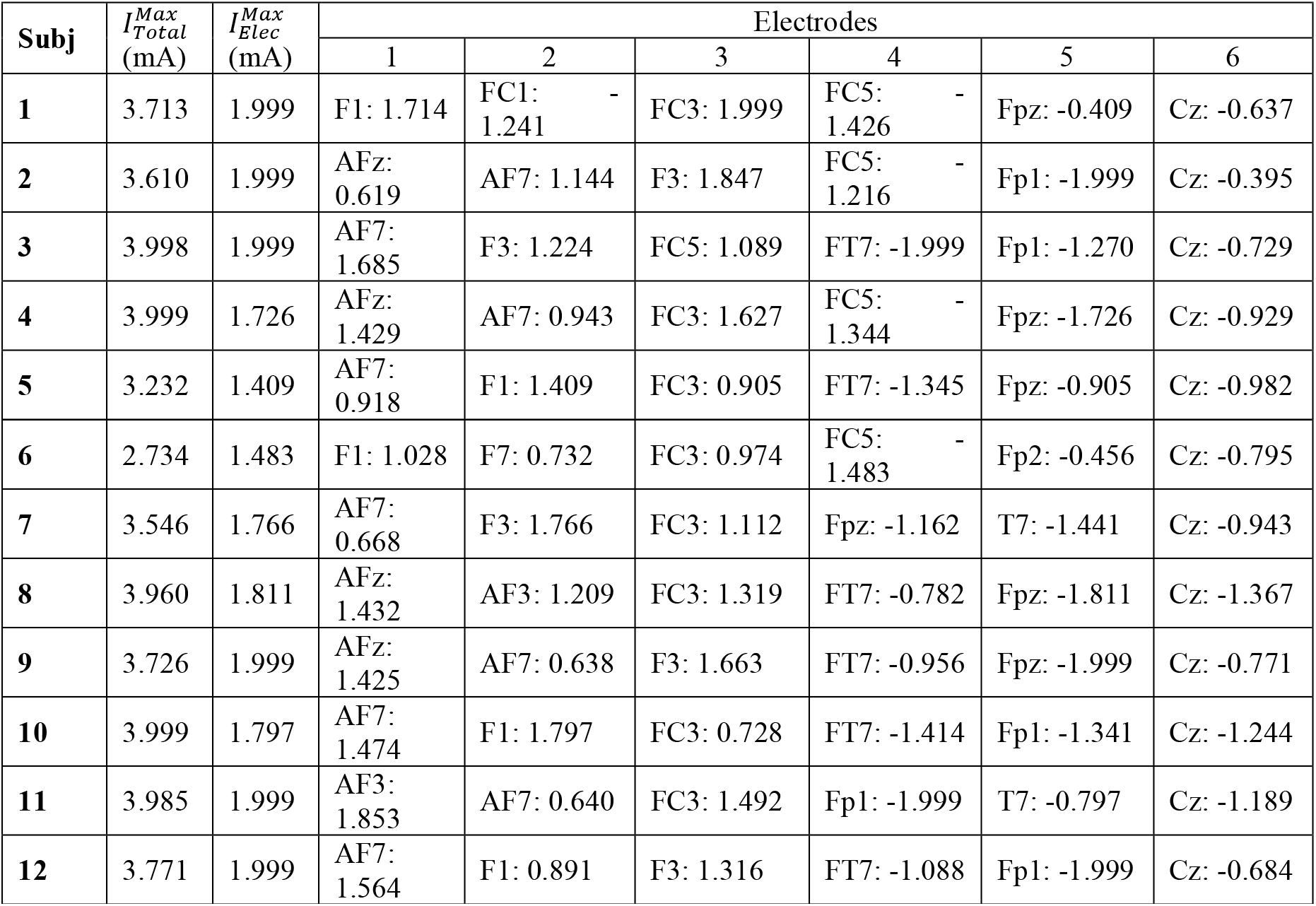
Results of the optimizations (electrode currents) for each of the subjects in this study. All currents in mA.

### 3. Impact of electrode modeling on results

The electrodes represented in the bipolar montages in this study are based on the circular SPONSTIM electrodes (https://www.neuroelectrics.com/products/electrodes/sponstim/). The latter consist of a conductive rubber circular pad (radius of 20 mm), connected to a 9.5 mm diameter electrode connector, sitting on top of the circular sponge electrodes (area of 25 cm^2^, radius of 28.2 mm). To determine the optimal way to model this electrode we compared the E-field distribution in models where the rubber sponge and metal connector were accurately represented, and in a model where the electrodes were modeled as homogeneous. ***Figure s1a*** compares both models of the electrodes. These calculations were performed in only one of the subjects of this study (subject 1). The rubber sponge was modeled as a material with a conductivity of 40 S/m (a value closed to what was measured in Fernandes et al., 2018), and the sponge underneath was modeled with the same value used in the homogeneous electrode model: 4.0 S/m. This resulted into some differences in E-field distribution under the electrodes with respect to the homogeneous model (see ***figure s1b***). However, these differences had a negligible effect on calculated E-field metrics such as the average E-field values: the relative difference (with respect to the values obtained with the complex electrode models) of the average E-field component values over both areas of interest were always less than 3% (see ***table s3***). The min/max values in the GM surface were also affected minimally by the type of electrode model.

These results show that the level of detail included in the electrode model has a negligible effect on the calculated average E-field metrics presented in this paper. As such, the results previously reported would not have changed much with the more accurate electrode model. These results would not necessarily apply if smaller region-of-interest areas had been considered, or for other electrodes geometries.

**Figure s1:**
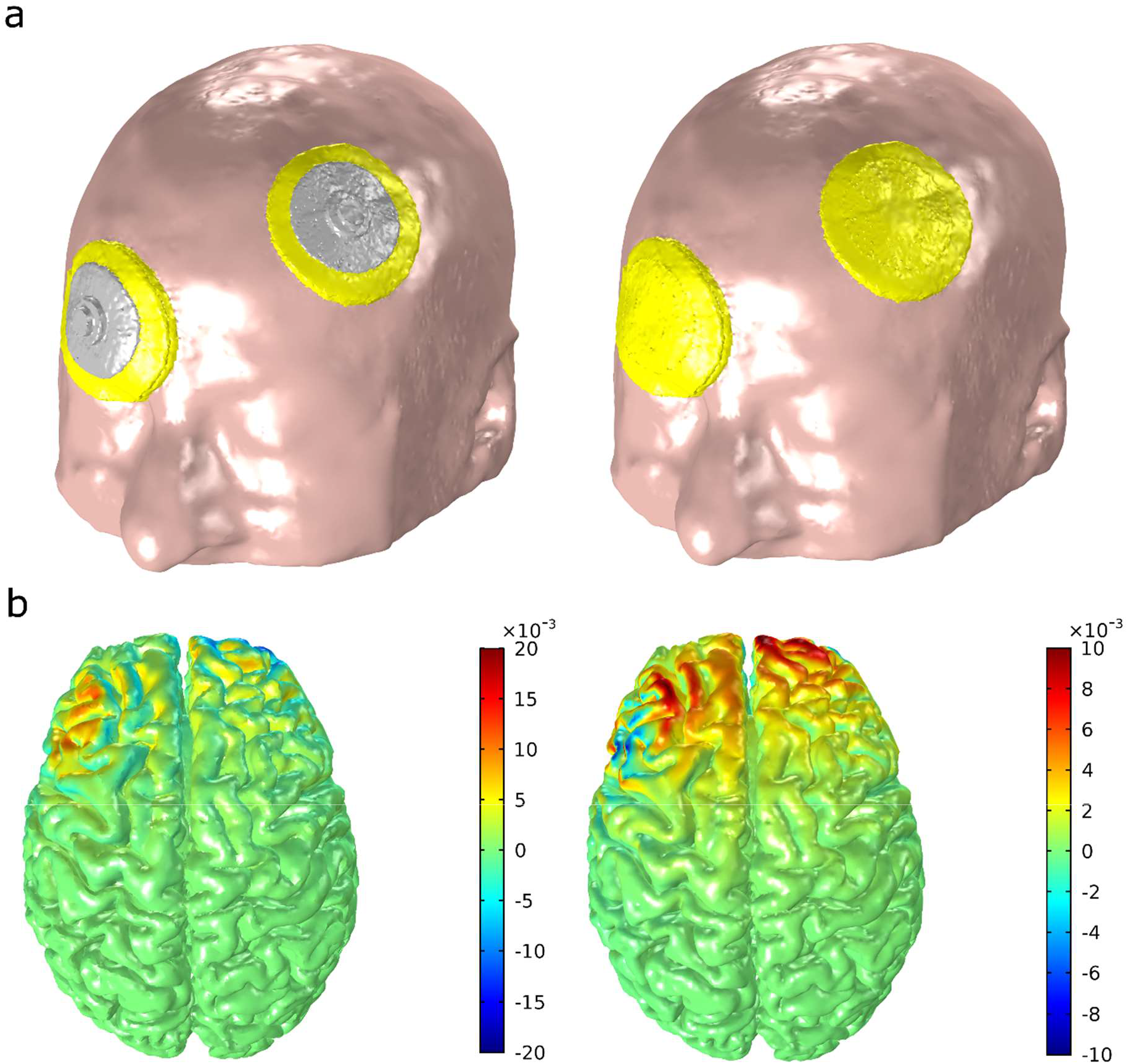
(a) Representation of the two electrode models compared in this study. (b) Absolute difference of the normal (left) and tangential (right) components of the E-field in the cortical surface between the homogeneous and rubber-pad electrode models (in V/m).

**Table s3:** Average values of the components of the E-field in the left DLPFC and right DLPFC for both electrode models. The max/min values of each component in the cortical surface are also shown. The % difference between the two values in presented in the last row: % *Δ* = (*Value_Homogeneous_* – *Value_Rubber pad_*)/ *Value_Rubber pad_* × 100.

|  | *Average* |  |  |  |  |  | *Min/max values* |  |  |  |
| --- | --- | --- | --- | --- | --- | --- | --- | --- | --- | --- |
| | $E_n$ (V/m) | | $E_t$ (V/m) | | $E$ (V/m) | | $E_n$ (V/m) | | $E_t$ (V/m) | $E$ (V/m) |
| *Model* | Left DLPFC | Right DLPFC | Left DLPFC | Right DLPFC | Left DLPFC | Right DLPFC | Max | Min | Max | Max |
| Homogeneous | 0.053 | -0.020 | 0.144 | 0.127 | 0.224 | 0.196 | 0.812 | -0.873 | 0.462 | 0.949 |
| Rubber pad | 0.052 | -0.020 | 0.148 | 0.130 | 0.229 | 0.200 | 0.812 | -0.875 | 0.463 | 0.950 |
| %$\Delta$ | 0.9 | -2.1 | -2.6 | -2.3 | -2.2 | -2.2 | 0.0 | -0.2 | -0.1 | -0.2 |

